# A novel *in vitro* assay for quantifying endothelial cell and pericyte interactions

**DOI:** 10.64898/2026.08.03.742618

**Authors:** David J Csordas, Lily C Cucuzzella, John S Kim, Shayn M Peirce

**Affiliations:** Department of Biomedical Engineering, University of Virginia; Charlottesville, VA, USA; Robert M. Berne Cardiovascular Research Center, University of Virginia; Charlottesville, VA, USA; Division of Pulmonary and Critical Care Medicine, University of Virginia; Charlottesville, VA, USA

**Keywords:** Endothelial cells, pericytes, microvasculature, microvascular stability, image analysis pipeline

## Abstract

**Objective:** Structural adaptations of capillary networks, through angiogenesis, arterialization, and regression, are implicated in many diseases, and gaining a better understanding of the cell-cell interactions that underpin these adaptations may lead to novel therapeutic discoveries for disease management. Endothelial cells and pericytes are the two cell types that comprise capillary networks. Experimental model systems have been developed to study the dynamic interactions between endothelial cells and pericytes, providing valuable insights into capillary development, cell-to-cell communication, and responses to growth factors and therapeutic agents.

**Methods:** In this study, we present a novel and simple co-culture system that uses commercially available primary human endothelial cells and pericytes, does not require microfluidic perfusion, and allows simultaneous observation of cell morphologies and interactions over time in 60 samples, enabling high-throughput analysis of multiple culture conditions with replicates.

**Results:** Image analysis pipelines were created to quantify microvascular adaptations, including one to measure colocalization between endothelial cells and pericytes, capturing dynamic coupling and uncoupling associated with capillary stability, angiogenesis, and regression. We validated the ability of our co-culture system to reproducibly represent the effects of fibrotic and angiogenic activation signals, including an FDA-approved drug, on endothelial cells, pericytes, and their coupling.

**Conclusion:** This novel, high-throughput microvascular screening assay enables quantification of microvascular dynamics in response to disease-relevant stimuli and therapeutics in a repeatable, real-time manner.

## INTRODUCTION

Capillary networks are composed of endothelial cells, which form the lining of the capillaries, and pericytes, which wrap around and support the capillary endothelial cells.^1^ Almost all acute and chronic diseases affect the local microvasculature. The impact of disease can lead to microvascular remodeling, changes in vascular permeability, regions of fibrosis, and increased immune trafficking.^2–5^ While some of these changes are compensatory, they can often contribute to disease severity and progression. Aberrant microvascular remodeling may contribute and propagate disease development and lead to its progression.^6^

Capillary adaptations are difficult to study *in vivo*. This is due to the nature of their incredibly small size, their ubiquity within and throughout tissues, and their heterogeneity between organs. Because of these difficulties, there is a need to model these interactions *in vitro* using human-derived biospecimens.

While there are a number of other models that attempt to recapitulate microvascular biology, they have shortcomings in that they may not use human-derived cells, may use other mesenchymal cell types rather than pericytes, use pericytes from the central nervous system to model peripheral networks, or require scientists to have access to flow systems that create a barrier to entry and limit the throughput of assays.^7,8^ Our goal for this study was to create a microvascular model system using readily available human-derived endothelial cells and peripheral pericytes. This system is scalable and can be performed at high throughput in 96-well plates.

## MATERIALS AND METHODS

### Cell Culture

Human umbilical vein endothelial cells (HUVECs) (Lonza, C-2519A) were cultured using standard protocols in microvascular endothelial cell growth medium-2 (EGM-2 MV) (Lonza, CC-3202). Human pericytes from placental origin (hPC-PLs) (PromoCell, C-12980) were cultured using standard protocols in pericyte growth medium-2 (PGM-2) (PromoCell, C-28041). Cells were maintained in flasks. For passaging, DPBS (Gibco, 14190-144) would be used to wash cell layers. Endothelial cells and pericytes were lifted off flasks using either 0.05% Trypsin-EDTA (Gibco, 25300-054) or StemPro Accutase (Gibco, A11105-01), respectively. Following centrifugation (200 x G, 5 minutes, room temperature), cells were replated or used in experiments. Both cell types were used in subsequent assays between passages 4 to 7.

### Live-Cell Staining

CellTracker Red (CTR) and Green (CTG) (ThermoFisher, C34552 and C7025) were used to label endothelial cells and pericytes, respectively. When cells reached 70% confluency, cell culture media was aspirated and replaced with either 25 µM CTR or 15 µM CTG in diluted in DMEM (Gibco, 11965-092) supplemented with 1% penicillin/streptomycin (P/S) (Gibco, 15140-122). For staining in a T75 flask, approximately 7.5 mL of media was used. Flasks were returned to the incubator (37°C, 5% CO_2_) for 30 minutes. The cell stain was subsequently aspirated and replaced with fresh culturing media. Cells were returned to the incubator overnight (or until treatment with mitomycin C). The next day (Day 0), media was aspirated, and cell fluorescence was confirmed via microscopy. All the following steps using these cells were performed in the dark in order to protect the fluorescent signal.

### Mitomycin C Treatment

This step was conducted on pericytes at least five hours after staining with CTG. Mitomycin C (Sigma, M5353) was diluted to 5 µg/mL in DMEM supplemented with 1% P/S. Cell culture media was aspirated from the flask, washed once with Dulbecco’s Phosphate Buffered Saline (DPBS) (Gibco, 14190-144), and diluted mitomycin C (approximately 8 mL in a T75 flask) was added. The flask was then returned to the incubator for 1 hour. After incubation, diluted mitomycin C was aspirated, and the flask was washed three times with DPBS. Finally, fresh cell culture media was added back into the flask, and the cells were returned to the incubator.

### Matrigel and Cell Preparation

Stock Matrigel (growth factor reduced, phenol red-free) (Corning, 35623) was thawed overnight at 4°C. After confirmation of live-cell staining, endothelial cells and pericytes were passaged using methods previously described. After centrifugation, both cell types were resuspended in EGM-2 MV and passed through a 100 µm cell filter (Falcon, 352360). Filtered cells were counted and resuspended to 5e6 endothelial cells/mL and 5e5 pericytes/mL. The cell suspension was kept on ice until the next steps.

On ice and using pre-chilled pipette tips, stock Matrigel was diluted to a working concentration of 5 mg/mL in EGM-2 MV. Endothelial cells and pericytes were then added and homogeneously mixed into the diluted Matrigel at a 5% volume for each cell type. For example, to make 1000 µL of cell-laden Matrigel, 50 µL of endothelial cells (5e6 cells/mL) and 50 µL of pericytes (5e5 cells/mL) were added to 900 µL of Matrigel (5 mg/mL). The final concentration of each cell type in cell-laden Matrigel is 2.5e5 endothelial cells/mL and 2.5e4 pericytes/mL.

Over ice, 40 µL of cell-laden Matrigel was added to each of the inner-60 wells of a pre-chilled 96-well plate (Corning, 3603). To reduce air bubbles, a negative-pipetting strategy was used to seed the gel. Prior to gelation, the well plate was tilted backward and gently tapped to ensure a level gel surface. DPBS (200 µL) was added to the outer wells to protect the experimental wells from edge effect evaporation. The plate was incubated for 30 minutes to allow the gel to fully form before adding subsequent media treatments. With this gel-volume added, each well of the plate had 10k endothelial cells and 1k pericytes. This 10:1 ratio was optimized for assay performance and is similar to ratios observed in healthy peripheral tissues ^9^.

Following incubation, 200 µL of EGM-2 MV was slowly added to each well and replaced back in the incubator to allow cells to adhere and begin to grow (Day 0, Hour 0). The following day, the media was aspirated and replaced with fresh media containing stimulation conditions.

### Stimulation and Treatment Preparation

ALK5 inhibitor (ALK5i) (Selleck, S8772) and Nintedanib (Selleck, S1010) were prepared to a stock concentration of 10 mM in 100% DMSO, then further diluted in EGM-2 MV to a 4x working solution. A matching dilution scheme was used to prepare DMSO (Invitrogen, D12345) for the control wells. TGF-β1 (R&D Systems, 7754-BH/CF) was reconstituted to 0.1 mg/mL in 4 mM hydrochloric acid (HCl). VEGF-A165 (R&D Systems, BT-VEGF) was reconstituted to 0.2 mg/mL in distilled water + 0.1% bovine serum albumin (BSA). TGF-β1 and VEGF-A were further diluted in EGM-2 MV to a 4x working solution. A similar dilution scheme was used to prepare compounds for two-day re-stimulation, but 2x solutions were prepared instead of 4x.

### Stimulation of Cells

Following aspiration of media after one day in culture, 100 µL of fresh EGM-2 MV was added to all wells, followed by 50 µL of 4x drug or 4x DMSO in the appropriate wells. The well plate was placed back into the incubator for 60 minutes for drug pre-treatment. Following this pre-treatment, 50 µL of 4x stimulant or EGM-2 MV was added to the appropriate wells, and the plate was returned to the incubator.

### Maintenance of Assay

Every two days in culture, a half-media swap was performed. To accomplish this, compounds were prepared at 2x working solution. 100 µL of spent media was removed, and fresh media was added back to the assay wells using a similar addition scheme as previously described to maintain a 200 µL media volume with appropriate compound concentrations.

### Microscopy and Live-Cell Tracking

All fluorescent and brightfield images were captured using a Leica THUNDER microscope (Leica, DMi8). A combination of 4x and 10x objectives (total magnification 40x and 100x, respectively) was used. For 10x objective images, five fields of view were randomly chosen in each well. Assay timing is anchored at the time of initial media addition as 0 hours. All wells were imaged at least once per day (e.g., at 24, 48, 72, and 96 hours). A subset of wells was chosen for time-lapse microscopy, during which images were collected every hour for 96 hours, with some gaps to allow for data saving and the capture of once-per-day images of other wells. The well plate was maintained in a stage-top incubator (Okolab, UNO-TH-CO2) that maintained environmental conditions (37°C, 5% CO_2_).

### Viability Measurement

Cell viability was quantified using a viability imaging kit containing Hoechst 33342 and propidium iodide (Invitrogen, R37610). Cell culture wells were set up in replicates. Each day, one set of replicates was incubated with media containing labeling stains for 30 minutes, then imaged. Remaining wells were maintained as normal, with half-media exchanges every other day. For quantification, nuclei were segmented into objects using StarDist in ImageJ ^10^, and CellProfiler was used to export total cell and dead cell counts. The latter was indicated by nuclei co-labeled with propidium iodide.

### Image Analysis

Images were analyzed using a custom pipeline implementing both ImageJ and CellProfiler. Fluorescent images were corrected in ImageJ to increase contrast and reduce background. Endothelial cells and pericytes were segmented into objects in CellProfiler and mapped for co-localization. To control for cell density variability, pixels of overlap between fluorescent channels were normalized to pixels of endothelial cells and pericytes. CellProfiler was also used to determine microvessel length and width. AngioTool 2.0 was used to quantify microvessel continuity and branchpoint metrics.^11^ Imaris was used to track cells, quantify distance, and measure velocity across time points.

### Statistical Analysis

When comparing two conditions, unpaired t-tests were used to test for significance. For more than two conditions, one-way ANOVAs with Sidak’s multiple comparison test or repeated-measures one-way ANOVA with Dunnett’s multiple comparisons test post-hoc analysis were used. GraphPad Prism was used for all graphs and statistical analysis.

## RESULTS

### Microvascular Assay Design and Optimization

Initial experiments were conducted to optimize gel type, cell concentration, media composition, cell seeding density, and endothelial-to-pericyte cell ratio. Primary readouts for these optimization assays were endothelial microvessel shape and quantity as a metric for healthy endothelial cells, and pericyte proliferation and morphological changes.

A schematic of the optimized assay setup is shown in **Figure 1**. One day prior to assay setup (Day −1), both cells are pre-labeled with CellTracker, and pericytes are treated with mitomycin C. Mitomycin treatment is performed 5 hours after CellTracker labeling. The CellTracker enables live-cell imaging and tracking throughout the experiment, while mitomycin temporarily arrests pericyte proliferation. This allows pericytes to interact with endothelial cell microvessels in culture without over proliferating the assay.

**Figure 1:**
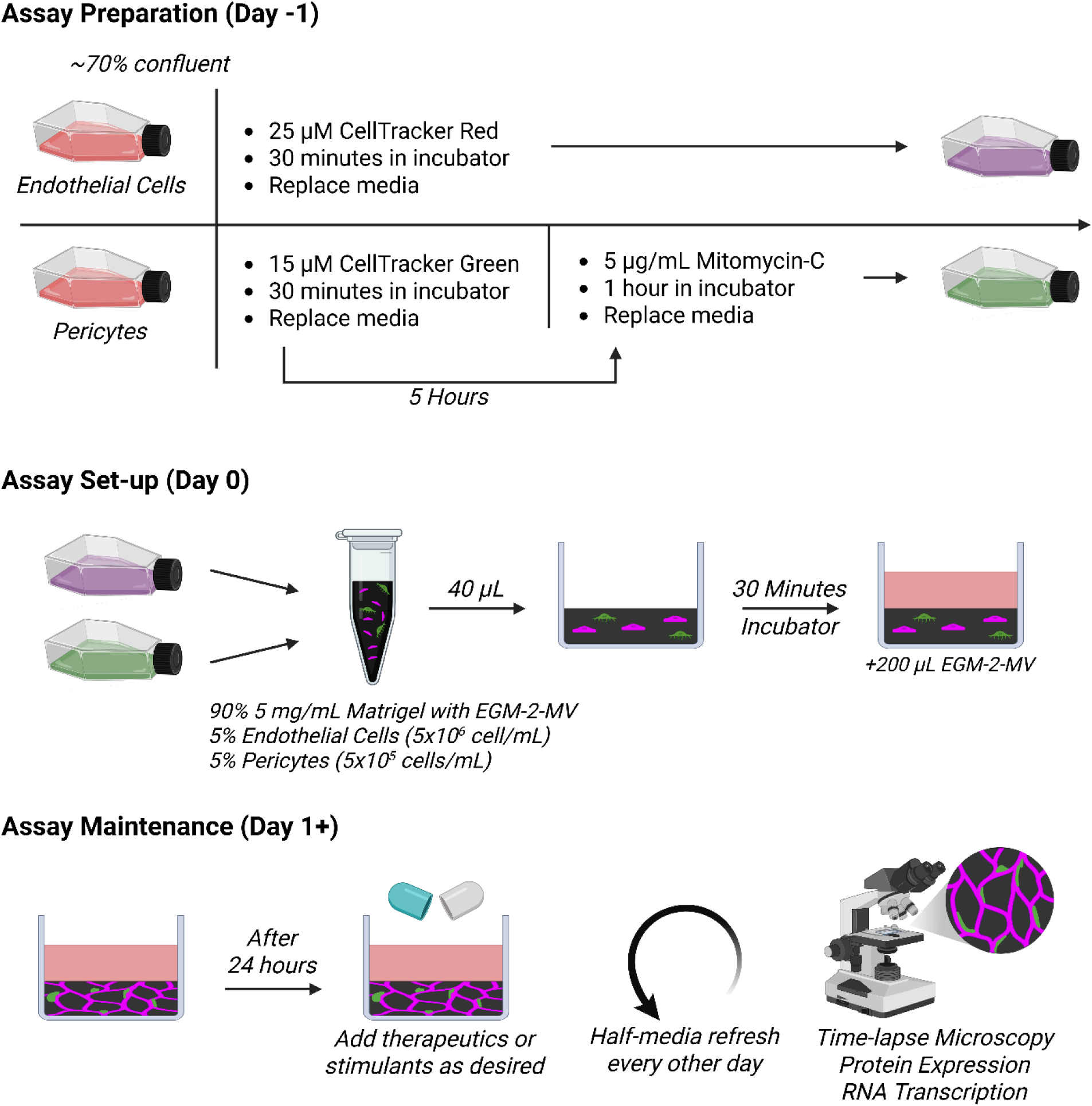
Summary of co-culture microvascular formation assay. Endothelial cells and pericytes are prepared in separate flasks prior to combination, embedding, and resuspension in Matrigel. Cell-laden Matrigel is seeded in 96-well plates, and microvessels begin to form overnight. Stimuli and compounds of interest can be added after 24 hours in culture, and the assay can be observed for seven days, with half-media exchanges every other day. At takedown, supernatant can be collected, and cells can be prepared for readout assays such as RNA transcriptome analysis, immunocytochemistry, or Western Blot.

Throughout the assay, cells maintained high viability (>94%) and were proliferative (**Fig. 2A-B**). Endothelial cells also formed multicellular vessel-like structures, with CD31 expressed at cellular junctions (**Fig. 2C**).

**Figure 2:**
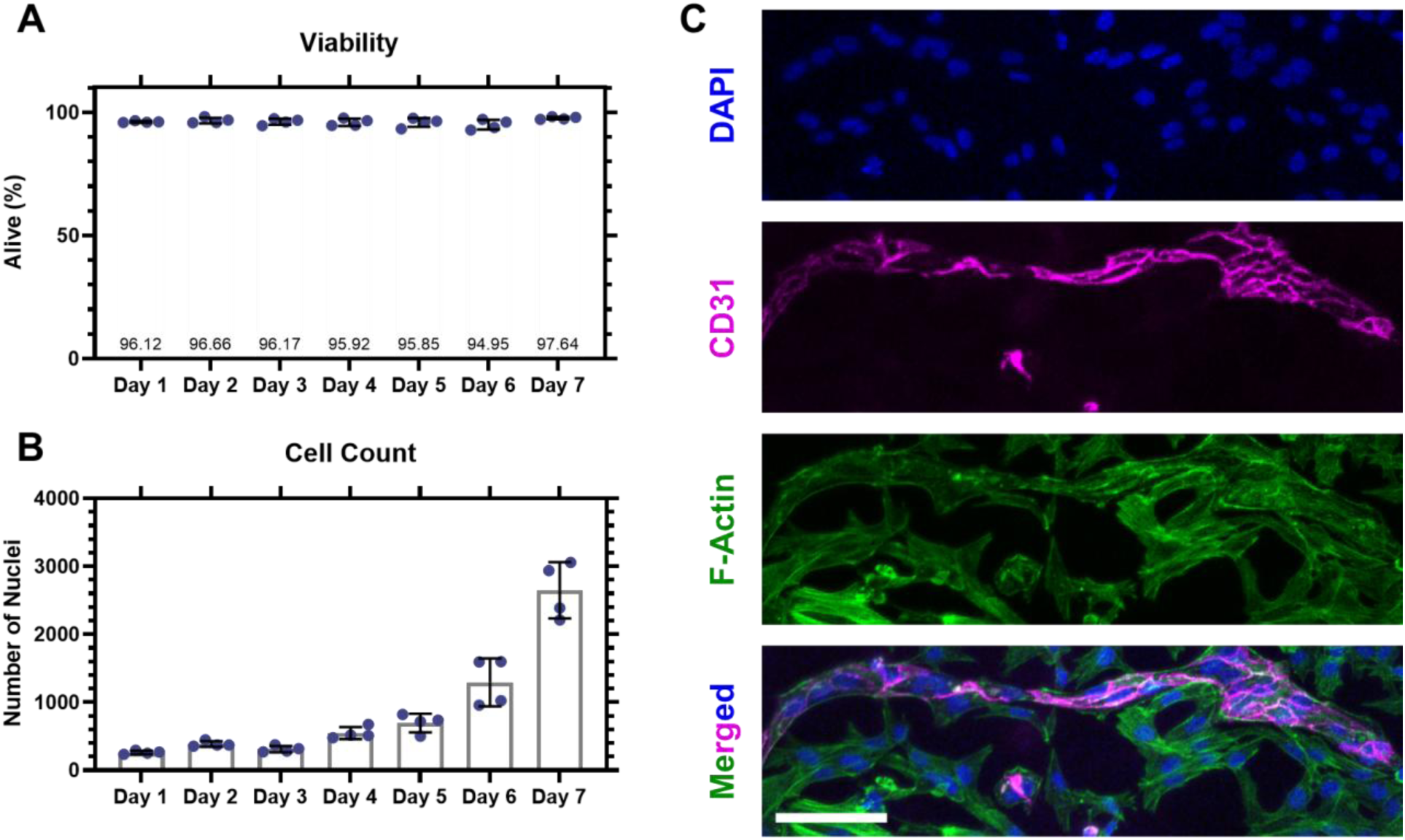
Endothelial cell and pericyte microvessel dynamics over time. **A)** Viability of cells across seven days in culture showed >94% viability on each day. **B)** Cell count on each day shows that cells are proliferating throughout the assay. Each point represents the average across five independent fields of view in each well (n = 4 wells per day). **C)** Immunofluorescence staining of cell nuclei (DAPI), endothelial cells (CD31), cytoskeletal component (F-Actin), and merged. Cells that are CD31-/F-Actin+ are pericytes. Scale bar represents 100 µm.

### Imaging and Quantifying Microvascular Structure and Dynamics Over Time

Changes in cell morphology over time can be compared across conditions (**Fig. 3A**). Compared with endothelial cells in monoculture, endothelial cells grown with pericytes are more numerous and cover a larger area (**Fig. 3B**). In co-culture, endothelial microvessel structures were significantly larger and longer than mono-culture controls. The width of each microvessel was similar in both cultures.

**Figure 3:**
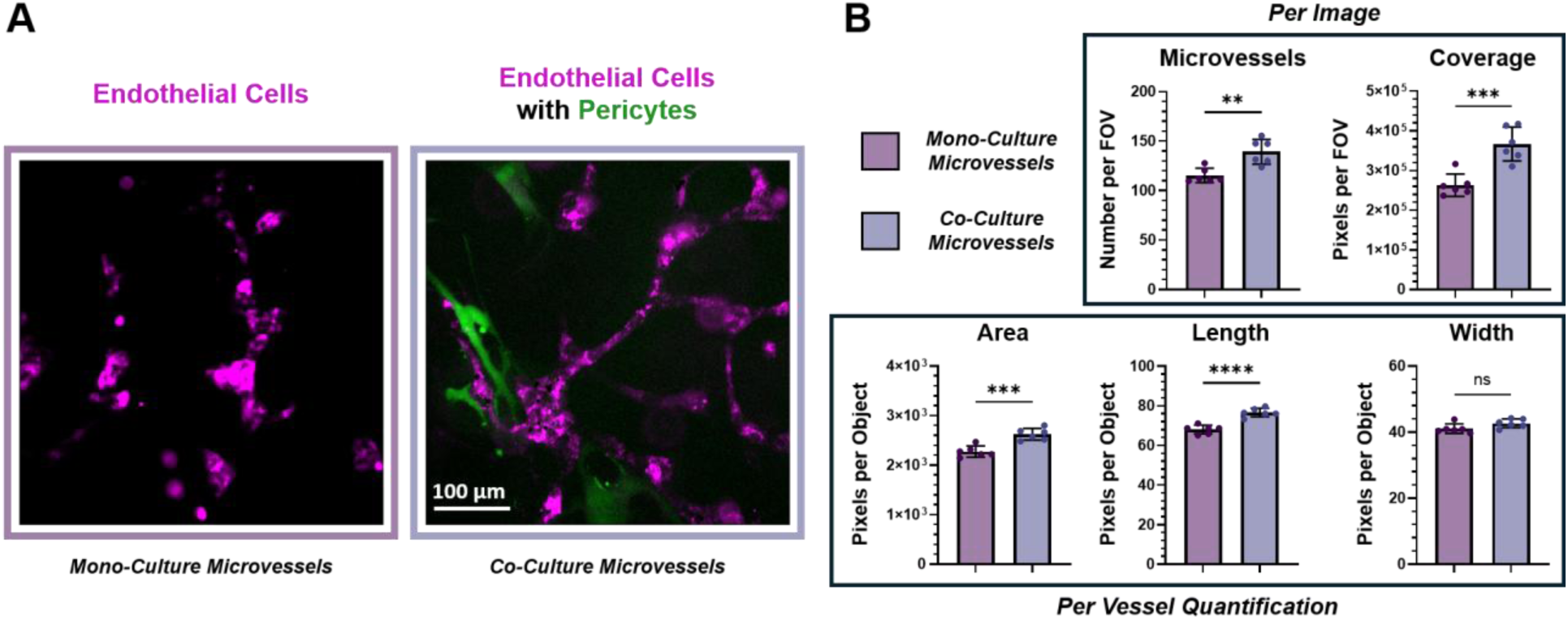
Microvascular morphological differences between cultures with and without pericytes. **A)** Mono-culture and co-culture with endothelial cells (magenta) and pericytes (green). Scale bar represents 100 µm. **B)** Endothelial microvessel characteristics quantified through a CellProfiler analysis pipeline. Data were either reported as per field of view (FOV) or per microvessel object across five independent fields of view in each well (n = 6 wells per condition). Images and microvessel metrics were quantified after four days in culture. Unpaired t-test used to quantify significance; ns=p>0.05, **=p<0.01, ***=p<0.001, ****=p<0.0001.

Time-lapse microscopy and individual cell tracking is possible when implementing a microscope-stage top incubator. Cell tracking can show microvessel shape dynamics, cell interactions, and division (**Fig. 4A**). Cell migration speed and distance can also be measured over time. Across a 24-hour observation window, when cells were treated with nintedanib, a triple receptor tyrosine kinase inhibitor targeting VEGFR, PDGFR, and FGFR, endothelial cells exhibited significantly decreased migration speed, shorter migration distance, and greater distance from neighboring cells compared to the DMSO control (**Fig. 4B**). The time during which cell tracks were identified by analysis software was not different between the two groups.

**Figure 4:**
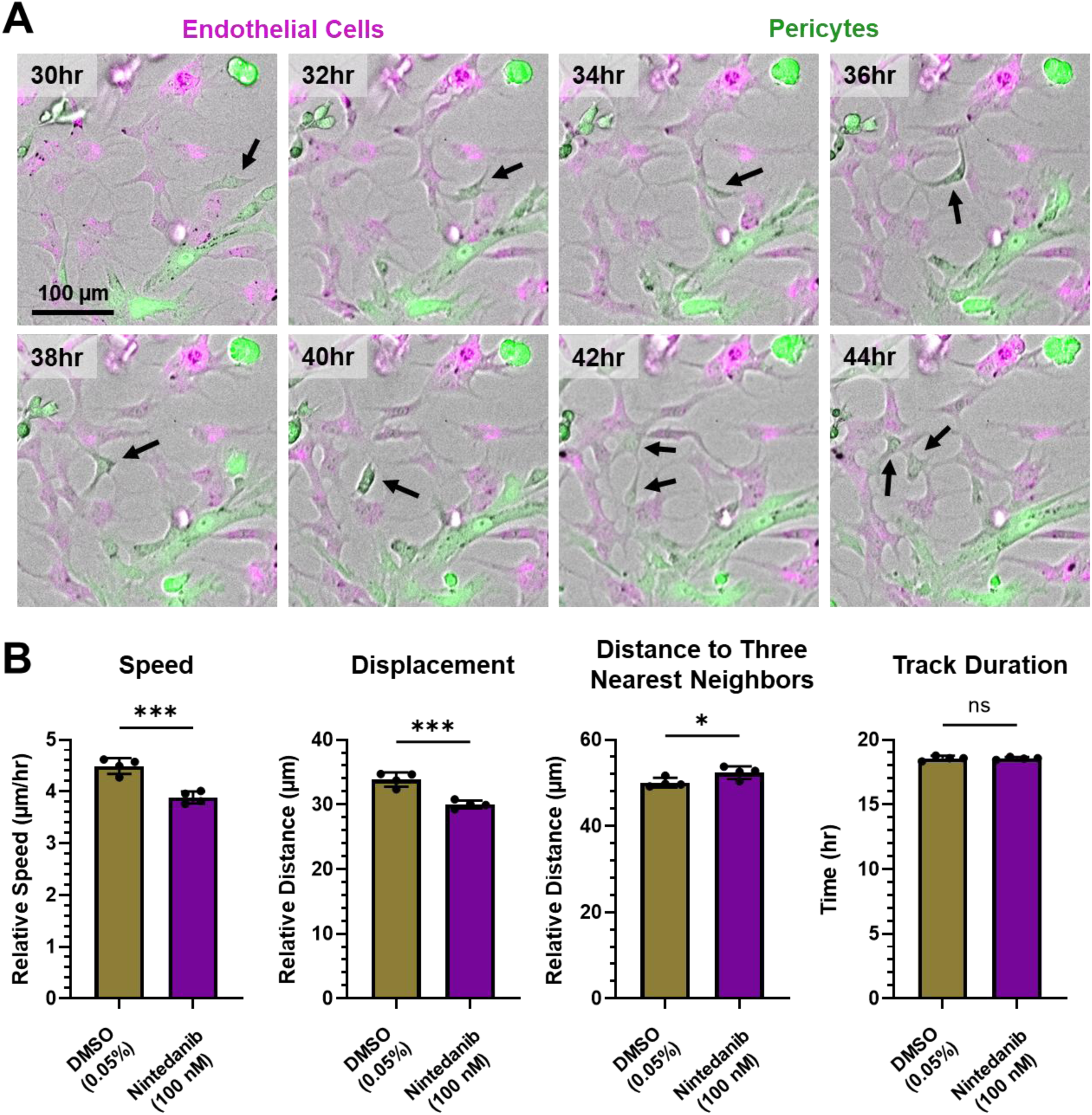
Live-cell tracking of microvascular cells. **A)** Time-lapse images of endothelial cells (magenta) and pericytes (green). Still images begin at hour 30 in culture and continue every two hours until hour 44. Scale bar represents 100 µm. **B)** Quantification of endothelial cell movement between hours 24 and 48 in culture, with images captured at one-hour intervals. Time-lapse images were analyzed using Imaris tracking software. Each point represents the average of nine independent fields of view per well (n = 6 wells per condition). Unpaired t-test used to quantify significance; ns=p>0.05, *=p<0.05, ***=p<0.001.

### Automated Measurement of Endothelial Cell and Pericyte Colocalization

The physical proximity, or coupling, between endothelial cells and pericytes leads to microvascular stabilization, maintenance of endothelial tight junctions, and inhibition of cellular migration.^12,13^ In homeostatic tissue, pericytes are coupled to endothelial cells and act as stabilizers to the microvascular unit. Increased colocalization reflects greater microvascular cell coupling and thus indicates a more mature and stable microvasculature. Decoupling of endothelial cells and pericytes occurs as part of the normal microvascular remodeling response in healthy tissue. However, in a pathological setting, this process can be perturbed, leading to aberrant microvascular structures characterized by impaired blood transport and poorly formed endothelial cell tight junctions ^14^.

To quantify colocalization in this assay, a custom image analysis pipeline was built to sort, correct, and identify cellular objects using live-cell tracking dyes (**Fig. 5A**). The ultimate output of the pipeline is the number of colocalized pixels, which counts the individual pixels of direct overlap between cell types (**Fig. 5B**). To correct for cell seeding and spreading variability between wells and days, the pixels of direct overlap are divided by the total number of endothelial cell and pericyte object pixels. The resulting output is normalized pixels of colocalization.

**Figure 5:**
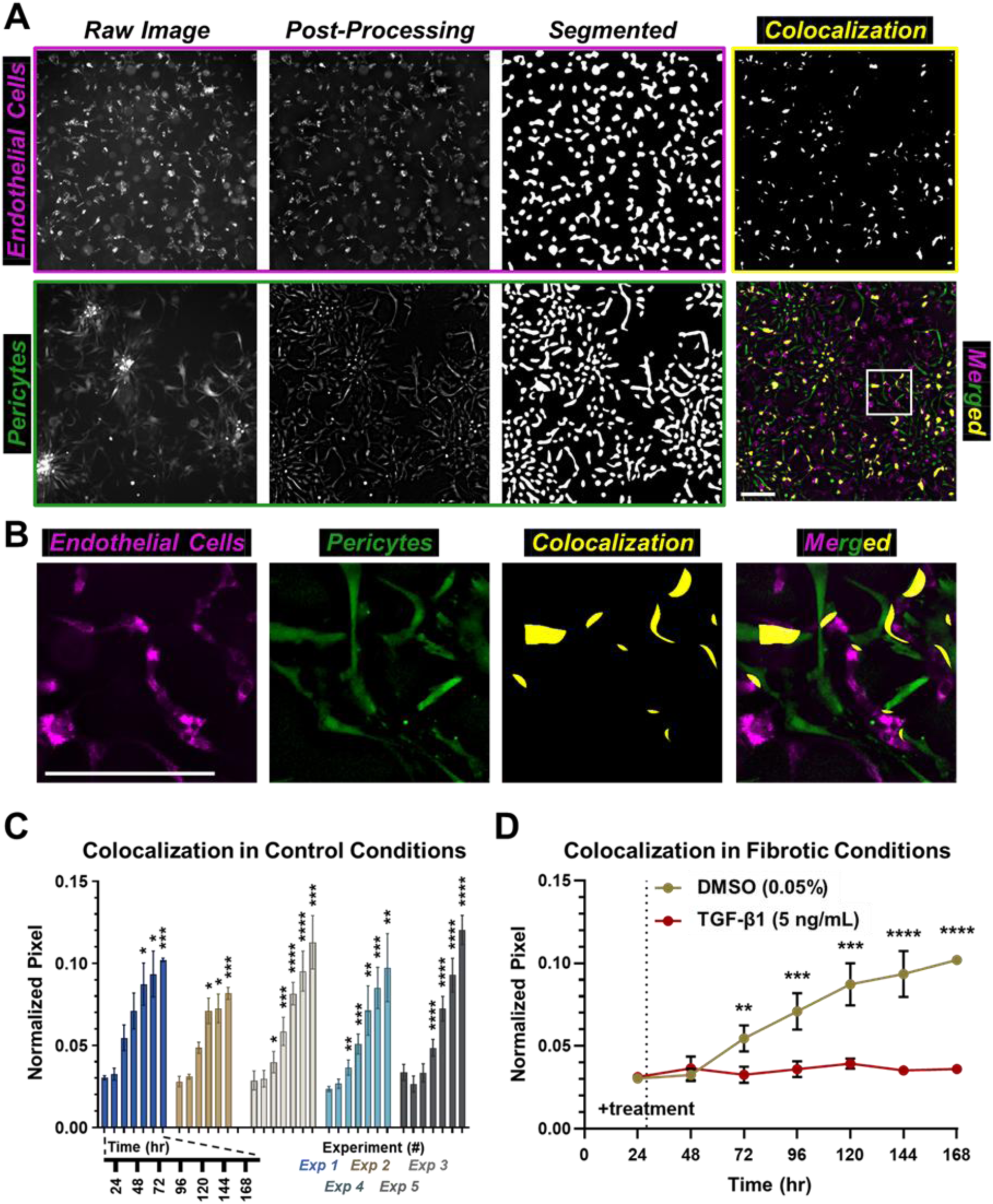
Pipeline to quantify colocalization between endothelial cells and pericytes. **A)** Images of fluorescently labeled cells are captured, filtered, and segmented. Direct colocalization is measured by calculating the overlapping pixels between endothelial cells and pericytes. **B)** Inset from **A** showing endothelial cells (magenta), pericytes (green), and area of colocalization (yellow). The scale bars for **A** and **B** are 200 μm. **C)** Colocalization was quantified every 24 hours for seven days across five experiments (Exp) in cells treated with 0.05% DMSO. For every condition, five independent fields of view were averaged to calculate colocalization for each independent well. Number of wells for each experiment is as follows: Exp 1, n = 3; Exp 2, n = 3; Exp 3, n = 6; Exp 4, n = 6; Exp 5, n = 6. Significance was determined within experiments using a repeated-measures one-way ANOVA with Dunnett’s multiple comparisons test comparing colocalization at each sequential time to what was measured at 24 hours. **D)** Repeated colocalization measurements comparing conditions treated with 5 ng/mL of TGF-β1 compared to 0.05% DMSO control. Five independent fields of view were averaged to calculate colocalization for each n (n = 6). Standard deviations are not shown if smaller than the symbols used in graphing. Unpaired t-tests were used to quantify significance at each time point. For significance, *=p<0.05, **=p<0.01, ***=p<0.001, ****=p<0.0001.

Colocalization was tracked in cultures treated with 0.05% DMSO for seven days, with measurements taken every 24 hours across five independent experiments (**Fig. 5C**). Consistent across all experiments, colocalization increased over time, becoming significantly different from that measured at 24 hours, generally by 96 hours. Treatment with 5 ng/mL TGF-β1 after 24 hours, to simulate fibrotic conditions of a wound, resulted in significantly less colocalization than in controls (**Fig. 5D**).

### Microvascular Responses to Growth Factors and Therapeutics

Exposure to pro-fibrotic (TGF-β1) and pro-angiogenic (VEGF-A165) factors modulated microvessel morphology. In cases, ALK5i, an inhibitor of the TGF-β1 activation cascade, and nintedanib, an inhibitor of VEGFR2, attenuated these morphological changes. TGF-β1 significantly decreased microvessel area and length, whereas VEGF-A had the opposite effect, increasing both metrics (**Fig. 6A-B**). TGF-β1 significantly increased microvessel diameter while VEGF-A165 did not induce a change (**Fig. 6C**). The average number of branch points, which is a measurement of microvascular bifurcation, was decreased by TGF-β1 and increased by VEGF-A165 (**Fig. 6D**). Taken together, VEGF-A165 induced a more migratory phenotype, with endothelial cells proliferating, stretching, and branching. In contrast, TGF-β1 induced regression and shrinkage of overall microvessel morphology, while leading to a widening of structures.

**Figure 6:**
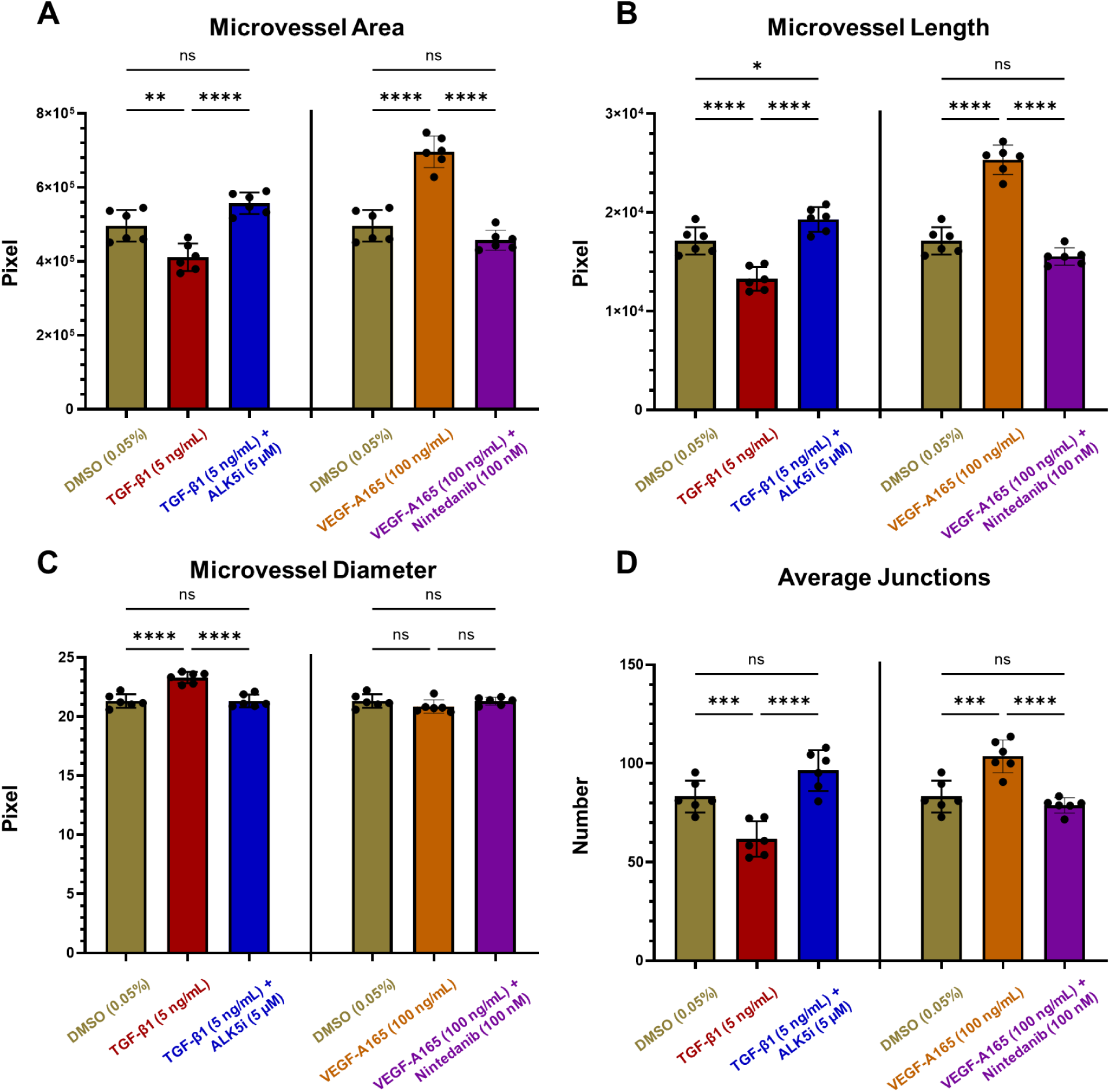
Changes to microvascular networks with exposure to growth factors. Microvessel **A)** area, **B)** length, **C)** diameter, and **D)** average number of junctions were quantified in conditions exposed to pro-fibrotic TGF-β1 (5 ng/mL) and pro-angiogenic VEGF-A165 (100 ng/mL). Respective inhibitors ALK5i (5 µM) and nintedanib (100 nM) were administered and compared to vehicle DMSO (0.05%) control. Five independent fields of view were averaged for each replicate (n = 6). AngioTool 2.0 was used to quantify microvessel attributes. One-way ANOVA with Sidak’s multiple comparison test was used to determine significance. For significance, ns=p>0.05, *=p<0.05, **=p<0.01, ***=p<0.001, ****=p<0.0001.

## DISCUSSION

### Challenges with Existing Co-Culture Systems and Opportunities for Improvement

Microvascular capillary networks are critical for proper tissue function, and injury can lead to local and systemic disease. It is critical to have models that can monitor capillary networks over time and in response to growth factors and therapeutics. Many variations of endothelial cell co-cultures exist that attempt to recapitulate capillary networks. Early models utilized a combination of endothelial cells and other cells, often fibroblasts, mesenchymal stem cells, or induced pluripotent stem cells.^15^ While these models can be robust and high-throughput, they fail to fully recapitulate the biological properties of pericytes. More complex derivations of these models involve incorporating flow cells into the tissue culture method.^16,17^ While these models produce phenotypes more similar to those of *in vivo* capillary systems, the use of flow cells can be a barrier for widespread adoption and severely limits the scalability of the assay for novel therapeutic testing.

More recent models utilize pericytes derived from central nervous system tissue (brain or retina).^18,19^ These systems can be built from human-derived materials, but are often made from a chimera of two species-derived cells cultured together.^13,20^ Though these systems are excellent for studying the microvascular system as it relates to the central nervous system, it is well established that pericytes derived from peripheral tissue exhibit distinct cell phenotypes and behaviors compared to those derived from central nervous tissue.^21^

Other papers have described systems utilizing human placental-derived pericytes, similar to the one described in this paper. These models benefit from the use of commercially available human-derived peripheral cells. However, current methods create co-culture systems that often collapse within 24 hours, making long-term analysis of microvascular remodeling in response to disease-relevant stimuli and treatments impossible.^22^

Given these challenges, the goal of this study was to create a high-throughput co-culture system using primary human-derived endothelial cells and peripheral pericytes to model capillary formation, stability, and dynamics over time.

### Design Constraints Driving Assay Development

Using primary human-derived cells and commercially available reagents, we aim to make this assay both translatable and broadly accessible. The assay should be stable and maintainable for at least one week to allow assessment of microvascular remodeling and molecular signaling dynamics can be accessed. It should also be designed to allow repeated measurements so that cell phenotypic changes in response to perturbations can be monitored in a high-throughput manner for testing of many conditions simultaneously. Finally, the goal of this work was to develop a relatively low-cost *in vitro* cell culture method using off-the-shelf materials and no additional equipment beyond what a laboratory would typically have.

To allow for long-term and stable culture, pericytes needed to be treated with mitomycin C to induce a temporary cell cycle arrest. This was necessary because in culture, pericytes will overgrow the endothelial cells. For media selection, while endothelial cells and pericytes were initially grown in separate media (EGM-2 MV and PGM-2, respectively), EGM-2 MV supported better microvessel formation when cells were combined in Matrigel. It was further determined that half-media exchanges every other day were optimal for microvessel formation. Despite perturbations with mitomycin and changes in media, the cells maintained high viability. Further, endothelial cells formed multicellular microvessel structures, and compared to endothelial mono-cultures, microvessels grown in the presence of pericytes are more abundant and longer in length.

To enable repeated measurements over time, cells were pre-labeled for live-cell imaging and tracking of morphological changes. Tests were conducted to ensure that the cell labels would remain stable throughout the assay and that the concentration used was not detrimental to cell viability. A secondary impact of allowing repeated measurements is that it enables more data to be collected from fewer samples, reducing the cost and experimental effort needed to conduct such studies.

This system was designed not to require flow to keep it low-cost, accessible, and high-throughput. While fluid flow is important for stimulating certain microvascular phenotypes, flow systems are a barrier to entry for many scientists and are typically low-throughput. By building our system to be compatible with a 96-well assay plate, we can test many conditions simultaneously with large numbers of replicates.

### Imaging Pipelines for Unbiased and High-Throughput Analysis

To take full advantage of the data collected throughout these experiments, we designed and implemented multiple image analysis pipelines to unbiasedly quantify cellular dynamics across thousands of images. One such pipeline **s**ought to quantify interactions between endothelial cells and pericytes by extracting colocalization information between the two cell types. In stable microvessels *in vivo*, we would expect pericytes to be colocalized and coupled with endothelial cells. We find that colocalization increases over time and can be modulated by pro-fibrotic stimuli.

To account for cells that are adjacent but not completely overlapping, the pipeline also outputs an additional characteristic called bloat colocalization. For this metric, the pipeline expands the identified cell object by five pixels in all directions, then measures pixel overlap. This number can be modified depending on the microscope magnification used to capture images. In all experimental tests, bloat colocalization yielded trends similar to those of direct colocalization.

To facilitate easy adoption of the methods described in this work, the image analysis pipelines developed to identify microvessel shape, structure, and cell colocalization will be made available on GitHub upon publication.

### Response to Stimuli and Therapeutic Intervention

The utility of this model system is to test disease-relevant stimuli and therapeutics on capillary networks in a high-throughput manner. Growth factors were used to assess microvascular responses to angiogenic and fibrotic conditions. Stimulation of angiogenic VEGF-A165 led to a pro-migratory phenotype, as endothelial cells proliferate and elongate. The addition of fibrotic TGF-β1 induced microvascular regression, as cells shortened and widened. This widening may indicate endothelial cells undergoing mesenchymal transition (EndoMT).

Therapeutics can also be tested in this system to better understand how they may impact microvascular dynamics. As expected, established compounds targeting both angiogenic and fibrotic activation pathways attenuated the morphologic changes induced by VEGF-A165 and TGF-β1. These inducible morphologic changes define the dynamic range of the culture that could be used in further studies to test novel therapeutics for anti-angiogenic and anti-fibrotic effects on capillary networks in a high-throughput manner.

Using time-lapse microscopy, cellular motility and migration speed can be tracked over time. We would expect that inhibiting growth factor receptors could modulate cell motility and movement. Indeed, compared to cells treated with nintedanib, an inhibitor of VEGFR, PDGFR, and FGFR, control endothelial cells migrated faster and moved in a more collective manner. Endothelial cells typically migrate up VEGF gradients, so if this cascade is blocked, it may blunt endothelial migration when it should otherwise occur.

This demonstrates how therapeutic compounds can be accessed for both their positive and negative effects on microvascular systems. While nintedanib is a potent antifibrotic and is approved for the treatment of pulmonary fibrosis,^23^ its mechanism of action may have off-target effects on the microvascular system.^24,25^ This assay allows scientists to assess these effects before moving to studies with animals or humans. By screening therapeutics, compounds can be derisked for microvascular impacts.

Disease-relevant stimuli can also be used to perturb the system. Beyond growth factor stimulation, cytokine cocktails or stimuli derived from patient samples, such as serum, could also be used. This would allow researchers to study, non-invasively, the effects of cytokine mixtures present in the disease microenvironment on microvascular systems. Novel therapeutics can also be assessed for their potential to stabilize microvasculature. Some research has proposed that YAP/TAZ inhibitors may act to stabilize microvascular abnormalities that occur during fibrotic disease.^26^ Other therapeutics, such as omega-3 supplementation, have been hypothesized as having vascular protective characteristics.^27^ This assay could be used to test these hypotheses directly and in a high-throughput manner.

## PRESPECTIVE

1. A novel, high-throughput microvascular screening assay using commercially available, primary human-derived endothelial cells and pericytes. This enables rapid and inexpensive hypothesis testing of microvascular systems.
2. Multiple computational analysis metrics and pipelines were created to unbiasedly quantify microvascular dynamics.
3. Biologically-relevant growth factor stimuli and inhibitors significantly altered microvascular dynamics and resulted in quantifiable changes in endothelial and pericyte interactions.
4. Future studies should interrogate the effect of novel therapeutics for the potential to stabilize microvasculature.

## STATEMENTS AND DECLARATIONS

### Data Availability

Data is available upon request

## Acknowledgments

Supported by National Heart, Lung, and Blood Institute (NHLBI) grants T32-HL007284 (DJC), K23-HL150301 (JSK), R01-HL176659 (JSK), and R01-HL155143 (SMP). Some figures featured in this manuscript were, in part or fully, created in https://BioRender.com.

